# Optimisation of lytic herpes simplex virus infection in human induced pluripotent stem cell derived cortical neurones

**DOI:** 10.1101/2025.08.22.671689

**Authors:** Daniel A. Nash, Alex S. Nicholson, Henry G. Barrow, Viv Connor, Colin M. Crump, Janet E. Deane, Stephen C. Graham

## Abstract

Herpes simplex virus (HSV)-1 infection of cortical neurones is a leading cause of encephalitis. While we have substantial knowledge about the molecular virology of HSV-1 lytic infection in cells of the periphery, like keratinocytes or fibroblasts, we know much less about infection of human neurones owing to the challenges of working with neuronal cell-based models. Here we demonstrate the use of a human induced pluripotent stem cell (iPSC)-derived cortical neurone model (i3Neurones) for HSV-1 infection. i3Neurones are highly scalable and can be rapidly and efficiently differentiated into an isogenic population of cortical glutamatergic neurones. We show that i3Neurones support the full HSV-1 lytic replication cycle. We present an optimised protocol for the infection of i3Neurones with HSV-1 that allows their synchronous infection at near-100% efficiency, and optimised fixation methods that preserves organelle and neurite structure for immunocytochemistry analysis. Our study highlights i3Neurones as a robust, scalable platform for microscopy and biochemical studies of HSV-1 and other neurotropic pathogens.

**Data summary:** The authors confirm all supporting data, code and protocols have been provided within the article or through supplementary data files.

## Introduction

Neuronal virus infections cause severe pathology. HSV-1 is the leading cause of viral encephalitis [1], causing 70% mortality in untreated patients and up to 19% in patients treated with antivirals, with survivors often suffering severe neurological sequelae [2]. Similarly, infection with enteroviruses such as EV-A71 cause encephalitis and acute flaccid paralysis [3]; Zika virus and Oropouche virus infection in adults can cause Guillain-Barré syndrome [4, 5]; and congenital Zika virus infection can cause microcephaly, decreased brain tissue, plus ocular and osteoskeletal abnormalities [6]. Furthermore, it is increasingly clear that infection with neurotropic viruses like HSV is a risk factor for developing common neurodegenerative diseases [7–9]. It is therefore important to identify robust and appropriate human cell-based systems to study the molecular basis of neuronal infection by HSV-1 and other neurotropic pathogens.

Multiple different systems have been used for the study of neuronal HSV-1 infection, especially in the context of latency (reviewed in [10]). *Ex vivo* infection of rat or mouse-derived ganglia that represent the natural site of HSV-1 latency are widely used [11–13], although the number of neurones that can be isolated even from a large number of animals is limited [14] and there are differences in interactions between HSV-1 and mouse versus human immune responses [15, 16]. *Ex vivo* studies of human neurones is possible using post-mortem specimens [17, 18], but availability and the capacity for genetic manipulation of the specimens is limited. Human embryonic or induced pluripotent stem cells (iPSCs) allow interrogation of infection following differentiation into neural stem cells, neurones or other glial cell types [19–23]. However, differentiation timescales can be long and the procedures labour-intensive. Human stem-cell derived organoids represent a powerful model for studying the functional consequences of HSV-1 lytic and latent infection in human neural stem cells, neurones and glia [24–28]. While excellent for transcriptomic analysis [25, 28], these models are not well suited to high resolution proteomics studies of infection such as quantitative temporal viromics [29], which require large numbers of homogenous cells (≥1×10^7^) and high levels of synchronous infection (≥90%) [30, 31].

Scalable cancer-derived neuroblastoma cell lines like SH-SY5Y are widely used for HSV-1 infection studies [21, 32, 33] but they have complex chromosomal aberrations [34, 35], are highly sensitive to the differentiation procedure used and yield mixed morphology populations [36]. Lund human mesencephalic (LUHMES) cells have been developed as models to study HSV-1 latency [37] and host shutoff during lytic infection [38]. Differentiated LUHMES resemble post-mitotic dopaminergic neurones [39] and they represent a powerful homogenous cell-based system for studies of neuronal infection. However, LUHMES cells are derived from the midbrain mesencephalon [40] whereas herpes simplex encephalitis (HSE) is generally localised to the temporal lobes [41]. As we know that innate immune programs differ between different classes of neurone [42], there is a need for additional scalable systems for the study of HSV-1 infection in the cerebral cortex.

Differentiation of human iPSCs via the expression of integrated transcription factors represents a promising approach to rapidly obtain isogenic populations of differentiated neuronal and other cell types [43]. For example, human iPSCs expressing the neuronal transcription factor Neurogenin3 (NGN3) [43] can be differentiated into sensory human neurones that have been used to characterise miRNAs and neuronal factors that regulate the efficiency of HSV-1 lytic replication or establishment of latency [44, 45]. Recently, a comprehensive analysis has shown that sensory neurones differentiated via NGN3 expression support synaptic firing plus lytic replication, latency and reactivation of HSV-1 [46]. However, given the clinical importance of HSE it is also necessary to have scalable, tractable systems to probe lytic infection of cortical neurones. This can be achieved via differentiation driven by Neurogenin2 [47]. Human iPSCs with doxycycline-inducible *Ngn2* in a safe-harbour locus [48] can be differentiated in 14 days into cortical glutamatergic neurones with close to 100% efficiency using a simple two-step protocol [49]. These integrated, inducible, and isogenic iPSCs (i3Neurones) exhibit robust synchronous neuronal firing [50] and are amenable to genetic manipulation [50, 51], making them suitable for precisely targeted functional studies. The ability to generate a homogenous isogenic population via activation of a stably-integrated master transcriptional regulator reduces experimental variability that can confound the interpretation of genome-or proteome-wide screening experiments [48], making these i3Neurones a robust platform for molecular discovery research.

Here we present optimisation and initial characterisation of lytic HSV-1 infection of i3Neurones, expanding the toolkit for biochemical and functional characterisation of neuronal HSV-1 infection.

## Methods

### Stem cell culture

Human fibroblast-derived iPSCs containing a doxycycline-inducible *Ngn2* transcription factor and an inactivated (dead) Cas9 gene in a safe-harbour locus [52] were provided by Michael Ward (National Institutes of Health, USA) and cultured as per [49]. Briefly, iPSCs were maintained in a 5% CO_2_ humidified atmosphere at 37°C in dishes pre-coated with hESC-qualified Matrigel (Corning 354277) diluted 1:100 in Dulbecco’s Modified Eagle Medium/Nutrient Mixture F-12 (DMEM/F-12; Gibco 11330032). Essential 8 medium (Gibco A1517001) was used for 24 hr culture and Essential 8 Flex medium (Gibco A2858501) for 72 hr culture, and colonies were subcultured by dissociation using 0.5 mM EDTA in PBS. iPSCs were dissociated to single-cell suspension with StemPro Accutase Cell Dissociation Reagent (Gibco A1110501) and seeded in medium supplemented with 50 nM Chroman1 (Rho-associated protein kinase (ROCK) inhibitor; Bio-Techne 7163/10).

### Neuronal differentiation

Differentiation of iPSCs into i3Neurones followed a two-step protocol of differentiation and maturation as outlined in [49]. In summary, 1.5–1.8×10^7^ iPSCs were seeded following Accutase dissociation in a Matrigel-coated 15 cm dish (Day 0) and incubated for three days in Induction Medium (IM): DMEM/F-12, supplemented with 1× N2 supplement (Gibco 17502048), 1× non-essential amino acids (NEAA, Gibco 11140050), 1× L-glutamine (Gibco 25030081), plus 2 μg/mL doxycycline (Sigma Aldrich D3072) to induce *Ngn2* expression. The medium was changed daily, being supplemented with 50 nM Chroman 1 for the first day of differentiation. At day 3 the i3Neurone precursor cells were dissociated with Accutase and frozen at −80°C in cryopreservation media comprising 90% (v/v) KnockOut Serum Replacement (Gibco 10828010) and 10% (v/v) DMSO before being stored in liquid nitrogen.

Day 3 i3Neurone precursor cells were cultured for a further 11 days (to day 14) in cortical neurone (CN) culture medium comprising Neurobasal Plus Medium (Gibco A3582901) supplemented with 1× B27 supplement (Gibco A3582901), 10 ng/mL Brain Derived Neurotrophic Factor (BDNF, PeproTech 450-02), 10 ng/mL Neurotrophin-3 (NT-3, PeproTech 450-03) and 1 μg/mL Laminin (Gibco 23017015). Frozen cells were thawed rapidly, diluted 10-fold in DMEM/F12 medium, pelleted by centrifugation (300 × g, 5 min), resuspended in CN medium supplemented with 2 μg/mL doxycycline and counted using a Countess II FL automated cell counter (Invitrogen). i3Neurones were seeded on plates coated with 100 μg/mL poly-L-ornithine (PLO) in half of the final culture volume of CN medium, incubated for 15 min at room temperature to allow cells to settle evenly, and then the remaining half of the medium was added before returning the cells to the incubator. When first seeding the cells the CN medium was supplemented with 2 μg/mL doxycycline, and half the volume of CN medium was replaced every three days until day 14, at which time the i3Neurones were used for experiments. At all stages, cells were cultured in a 5% CO_2_ humidified atmosphere at 37°C, and final differentiated neurones were confirmed as being free of mycoplasma.

### Non-neuronal cell culture

Vero (ATCC CRL-1586), U2OS (ATCC HTB-96) and U2OS pUL21-BirA*-HA cells (see below) were grown in DMEM supplemented with 10% (v/v) heat-inactivated foetal bovine serum (FBS) and 2 mM L-glutamine (complete DMEM) in a 5% CO_2_ humidified atmosphere at 37°C. All cells were frequently tested for mycoplasma and confirmed as mycoplasma-free.

U2OS cells expressing pUL21 tagged with an abortive biotin ligase (BirA*) [53] were generated by co-transfection of Flp-In T-REx U2OS cells provided by Gopal Sapkota (University of Dundee, UK) [54] with pOG44 (Invitrogen) and pcDNA5/FRT/TO (Invitrogen) encoding codon-optimised HSV-1 pUL21 (UniProt F8RG07) [55] with a C-terminal BirA* plus HA epitope tag. At 72 h post-transfection the culture medium was replaced with fresh medium containing 200 μg/mL hygromycin B and 3 μg/mL blasticidin. Selection of hygromycin and blasticidin resistant cells was allowed to proceed for 19 days, with medium being refreshed every 2-3 days as required. pUL21-BirA*-HA expression was induced by addition of 2 µg/mL doxycycline 24 h prior to use.

### Antibodies

Antibodies were used for immunofluorescence microscopy (IF), immunoblotting and virus neutralisation. See Table 3.1 for full details of antibodies and dilutions used.

**Table 1.**
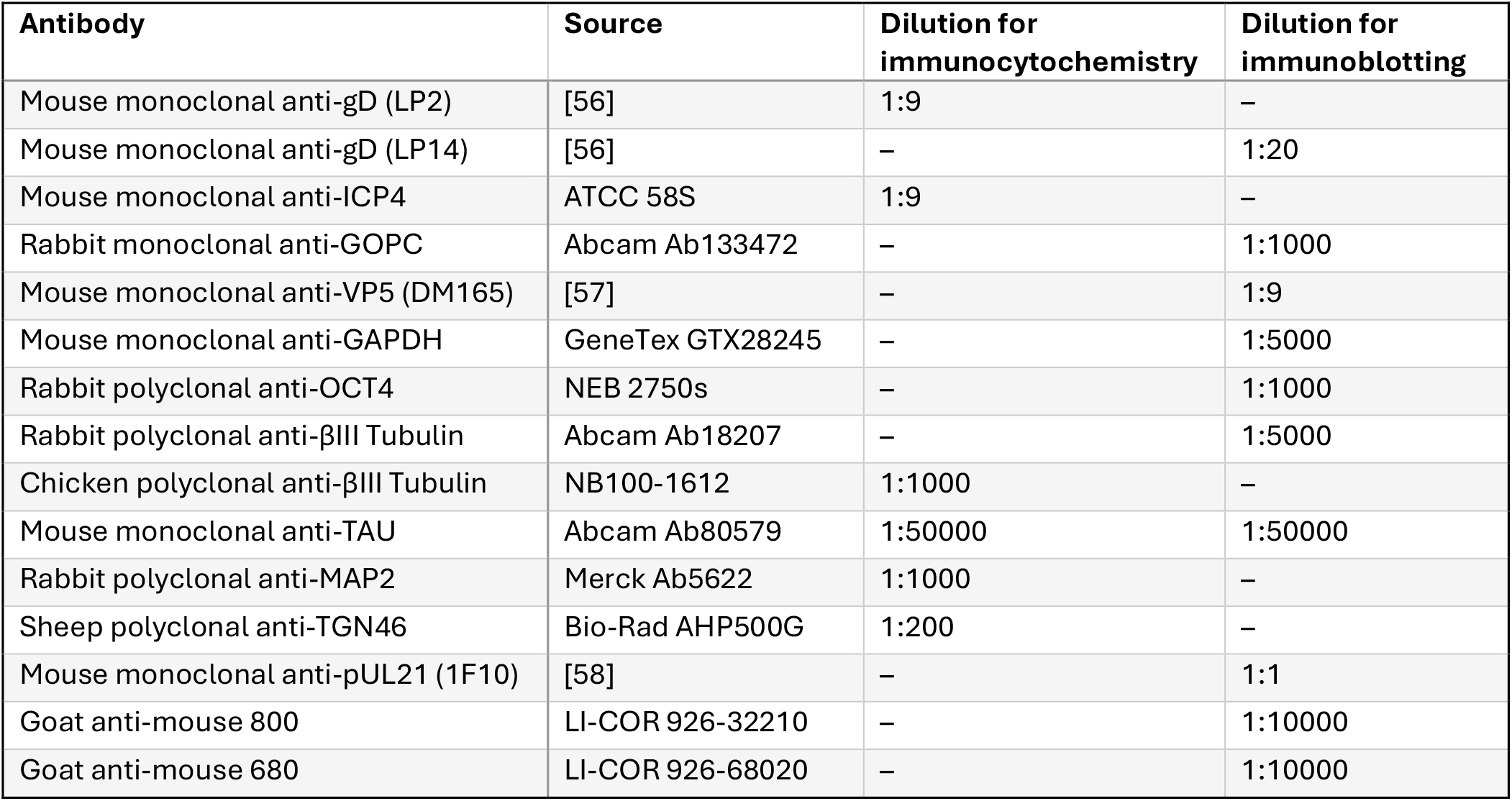
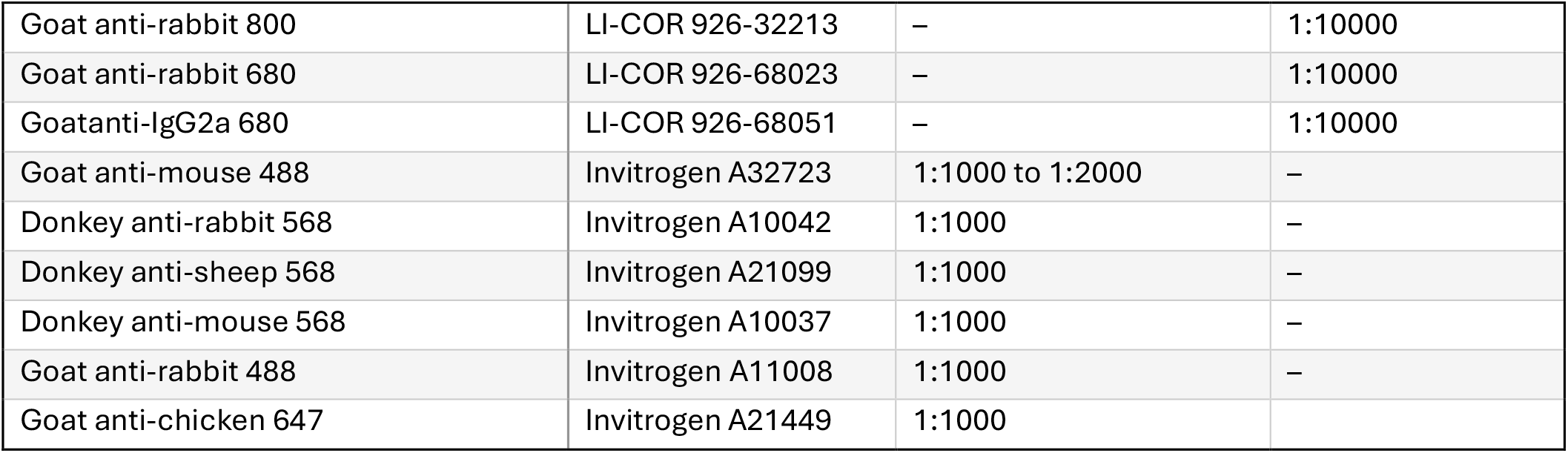
Antibodies used in this study.

### Viruses

Wild-type HSV-1 strain KOS (WT HSV-1) was derived from a bacterial artificial chromosome (BAC) encoding the KOS genome [59], as was wild-type HSV-1 with EFYP-tagged ICP0 and mCherry-tagged gC (WT timestamp HSV-1) [60]. A mutant timestamp HSV-1 lacking expression of pUL21 (ΔpUL21 timestamp HSV-1) was generated by two-step red recombination [59] of the WT timestamp BAC, introducing three stop codons into the pUL21 gene as described in [61]. Following the initial transfection of purified BAC DNA and pGS403 (encoding Cre recombinase) into Vero cells, all subsequent propagation was performed in complementing U2OS pUL21-BirA*-HA cells to minimise the chance of virus adaptation [58]. HSV-1 strain 17 was from Stacey Efstathiou (University of Cambridge, UK) and HSV-1 SC16 was from Tony Minson (University of Cambridge, UK).

Virus stocks were grown by infection of U2OS pUL21-BirA*-HA cells (ΔpUL21 timestamp HSV-1) or Vero cells (all others) at low (0.01) multiplicity of infection (MOI) for 3–5 days, until widespread cytopathic effect was evident. For timestamp viruses, cells were scraped into medium, freeze-thawed and sonicated at 50% amplitude for 40 seconds in a cuphorn sonicator before being clarified by centrifugation at 3,200×g for 5 min in a benchtop centrifuge. For all other viruses, the culture medium was supplemented with 0.5 M NaCl and 100 μg/mL dextran sulfate (7–20 kDa; SigmaAldrich 51227). The following day the supernatant was harvested, filtered through a 0.8 μm cellulose nitrate membrane (Nalgene 450-0080), virions were pelleted by centrifugation at 17,000 rpm in a Type 19 rotor for 45 min at 4°C, and viruses were resuspended in PBS with 10% (v/v) glycerol. For all, virus stocks were aliquoted and stored at −70°C until required.

### Virus titration

Samples serially diluted in 500 µL complete DMEM were used to infect 6-well plates containing confluent monolayers of Vero cells for 1 h at 37°C before being overlaid with 2 mL complete DMEM containing 0.3% high viscosity carboxymethyl cellulose (CMC) and 0.3% low viscosity CMC. After 3 days, cells were fixed with 3.7% (v/v) formal saline for 20 min before either being imaged using an Incucyte SX5 (Sartorius) with a 20× long working distance objective (NA 0.45) for fluorescent (timestamp) viruses or stained with 0.1% toluidine blue. In both cases plaques were counted manually.

### Virus infection

For neuronal infections, day 3 i3Neurones were seeded at specified densities and allowed to mmature. After maturation (day 14), conditioned medium was reserved, i3Neurones were washed with PBS and then infected with the indicated viruses in fresh CN medium at the relevant MOI. The time of virus inoculation was assigned 0 h post infection (hpi). Inoculation volumes for neurones per well were as follows: 100 µL for 96-well plates, 300 µL for 24-well plates, 600 µL for 6-well plates. Inoculated neurones were placed on a rocking platform at 37°C in a humidified 5% CO_2_ atmosphere. After 1 or 2 h incubation (as indicated), the inoculum was removed, cells were gently washed twice with PBS (unless stated otherwise), and washed cells were overlayed with a 1:1 mix of fresh and conditioned CN medium that had been clarified by centrifugation. Final volumes of overlay per well were as follows: 200 µL for 96-well plates, 1 mL for 24-well plates and 2.5 mL for 6-well plates. Vero cells were infected as above but using complete DMEM in place of CN medium.

### Immunoblotting

Day 3 i3Neurones were seeded at 2×10^6^ cells per well in a PLO-treated 6-well plate and allowed to mature. Where indicated, were infected at day 13 cells with MOI 5 for 1 h with HSV-1 strains KOS, strain 17 or SC16, or mock infected. For all, at day 14 cells plate were washed once with room temperature (RT) PBS and then ice-cold PBS containing 1% EDTA-free protease inhibitor cocktail (SigmaAldrich P8849) was used to detach the neurone layer from the well, fully suspending it by quickly and repeatedly dispensing down the edge of the well. The cell suspension was transferred to a microcentrifuge tube and pelleted (5000 × g, 5 min, 4°C) to remove the supernatant before lysis for 5 min using ice-cold RIPA buffer (50 mM Tris pH 8.0, 150 mM NaCl, 1 mM EDTA, 1% Triton X-100, 0.1% SDS, 1% EDTA-free protease inhibitor cocktail) and centrifugation (9000 × g, 10 min, 4°C) to remove debris. Undifferentiated i3Neurone iPSCs were grown to ~80% confluence in Matrigel-coated 6-well plates before being washed with PBS, lysed in situ, transferred to microcentrifuge tubes and the lysate clarified as above. For both, protein concentrations were analysed by BCA assay (Pierce) and normalised before samples were boiled (95°C, 5 min) in Laemmli sample buffer and separated by SDS-PAGE. Separated proteins were transferred to Protran 0.45 µm nitrocellulose membranes (Cytiva) using the Mini-PROTEAN system (Bio-Rad). Membranes were blocked using Tris-buffered saline (TBS; 50 mM Tris pH 7.6, 150 mM NaCl) supplemented with 5% (w/v) skim milk powder. Primary and secondary antibody incubations were performed in TBS supplemented with 0.1% TWEEN (TBS-T) and 5% (w/v) skim milk powder. Immunoblots were imaged using an Odyssey CLx (LI-COR) and analysed using Image Studio Lite (LI-COR).

### Immunocytochemistry

For confocal imaging, 13 mm #1.5 borosilicate glass coverslips were etched with 1 M nitric acid for 24 h before washing with ethanol and sterile PBS then coated with 100 μg/ml PLO for 24 h. Day 3 i3Neurones were seeded on prepared coverslips at a density of 1×10^5^ cells per coverslip and were allowed to mature. After maturation (day 14), cells were washed with PBS and fixed with the 4% (v/v) EM-grade formaldehyde (Polysciences) in a 250 mM HEPES pH 7.5 buffer on ice for 10 min followed by an 8% (v/v) formaldehyde solution in 250 mM HEPES pH 7.5 buffer at room temperature (RT) for 20 min, unless otherwise stated. Fixed i3Neurones were washed thrice with PBS and permeabilised at RT on a rocking platform using PBS supplemented with 0.2% Triton X-100 for 5 min or 0.1% saponin for 15 min. Cells were blocked for 30 min at RT using blocking buffer comprising PBS supplemented with 5% (v/v) FBS alone (Triton X-100 permeabilisation) or 5% (v/v) FBS plus 0.01% saponin (saponin permeabilisation) and incubated with primary antibody in blocking buffer for 2 hr at RT. Coverslips were washed in blocking buffer, incubated with secondary antibodies in blocking buffer for 1 hr at RT in the dark, then washed in blocking buffer, PBS and then MQW. Cells were mounted on microscope slides using Mowiol 4-88 (Merck) supplemented with 200 nM 4′,6-diamidino-2-phenylindole (DAPI) and left to dry overnight before being stored at 4°C. Slides were imaged using an EVOS M5000 (Invitrogen) using a 20× plan fluorite long working distance objective (NA 0.45) or a Zeiss LSM700 confocal laser scanning microscopy system mounted on an AxioObserver.Z1 inverted microscope with a 64× plan apochromat objective (NA 1.4).

For monitoring percentage infection, cells grown in PLO-treated culture vessels were fixed in situ using 4% then 8% formaldehyde in HEPES buffer and stained for ICP4 as detailed above. Immunostained cells were overlayed with propidium iodide (2.5 µg/mL in PBS + 0.02% sodium azide) to stain the nuclei and were imaged via phase contrast and fluorescence using an Incucyte SX5 (Sartorius) with a 20× long working distance objective (NA 0.45). Images were quantified using the Basic Analyser algorithm in the Incucyte analysis software (Sartorius).

For all, figures were generated using ImageJ [62, 63].

### Inoculation condition optimisation

Day 3 i3Neurones were seeded at 3×10^5^ cells per well in a PLO-treated 24-well plate and allowed to mature. Infection proceeded as described above with changes to the inoculation conditions as follows: cells were inoculated for 60, 90 or 120 min being rocked either manually every 15 min or automatically at a low speed on a rocking platform. The inoculum was removed and cells gently washed with PBS twice (unless stated otherwise), overlayed with a 1:1 mix of fresh and clarified conditioned CN medium. Cells were fixed at 16 hpi and percentage infection was monitored by immunocytochemistry using 4 to 8% formaldehyde in HEPES buffer for fixation and Triton X-100 for permeabilisation as described above. Data were analysed using a two-way ANOVA and significance of differences to 60 min sample was assessed using Sidak’s test in Prism 7 (GraphPad). The equivalency of variance across all data points at MOI 5 for the automatic versus manual rocking was assessed using Levine’s test [64] in Prism 7 (GraphPad).

### Percentage of infected i3Neurones at different MOIs

Day 3 i3Neurones were seeded at 3×10^5^ cells per well in a PLO-treated 24-well plate and allowed to mature. At day 13, Vero cells were seeded separately at 5×10^5^ cells per well. The following day, both i3Neurones and Vero cells were infected with a 2-fold serial dilution of WT HSV-1, with MOIs ranging from 10 to 0.0098. After 2 h the inoculum was removed and the cells gently washed twice with PBS (unless stated otherwise) before being overlayed with a 1:1 mix of fresh and clarified conditioned media. Cells were fixed at 16 hpi and percentage infection was monitored by immunocytochemistry as described above.

### Neurone survival following infection

Day 3 i3Neurones were seeded at 3×10^5^ cells per well in a PLO-treated 24-well plate and allowed to mature before being infected at MOI 5 for 1 h with HSV-1 strains KOS, strain 17 or SC16, or mock infected. Cells were washed twice with PBS, and then overlayed with a 1:1 mix of fresh and clarified conditioned CN medium supplemented with 2.5 µg/mL propidium iodide. Cell viability, defined as exclusion of the propidium iodide, was monitored by capturing phase contrast and orange fluorescence images every 6 h using an Incucyte SX5 (Sartorius). Images were analysed using the Basic Analyser algorithm in the Incucyte analysis software (Sartorius). Half the volume of CN medium, supplemented with fresh 2.5 µg/mL propidium iodide, was replaced every 3 days.

### Inoculum inactivation

Day 3 i3Neurones were seeded at 3×10^5^ cells per well in a PLO-treated 24-well plate and allowed to mature. The i3Neurones were infected at MOI 5 for 1 hr before the inoculum was removed and cells were either: washed thrice immediately with PBS; incubated with 25 μg/mL LP2 for 15 min at 37°C before three PBS washes; or washed once with citric acid (40 mM citric acid pH 3.0, 135 mM NaCl, 10 mM KCl) for 1 min before three PBS washes. Cells were overlayed with a 1:1 mix of fresh and clarified conditioned CN medium. At 3 and 24 hpi, neurones were harvested by thrice freezing the plate at −70°C and then thawing. Lysed cells were scraped into the medium and the virus concentration was assessed by plaque assay. Data were analysed using a one-way ANOVA and significance was assessed using Tukey’s test in Prism 7 (GraphPad).

### Fixative Condition Tests

Day 3 i3Neurones were seeded at 1×10^5^ cells per well on prepared coverslips in 24-well plates and allowed to mature. The i3Neurones were infected with WT HSV-1 at MOI 3. At 16 hpi, neurones were washed with PBS and fixed using one of the following four treatments: 4% (v/v) EM-grade formaldehyde in a 250 mM HEPES pH 7.5 on ice for 10 min then an 8% (v/v) formaldehyde solution in 250 mM HEPES pH 7.5 at RT for 20 min; cytoskeletal fixing buffer (300 mM NaCl, 10 mM EDTA, 10 mM glucose, 10 mM MgCl_2_, 20 mM PIPES pH 6.8, 2% sucrose, 4% formaldehyde) on ice for 15 min; 100% methanol on ice for 10 min; or glyoxal buffer pH 4 (3% (v/v) glyoxal, 20% (v/v) ethanol, 0.75% acetic acid, pH adjusted with NaOH) on ice for 30 min followed by a further 30 min at RT. Fixed neurones were washed, permeabilised, stained and imaged as described above.

### HSV-1 spread assay

Day 3 i3Neurones were seeded at 3×10^5^ cells per well in a PLO-treated 24-well plate and allowed to mature before being infected at MOI 0.1 for 2 h with timestamp viruses, washed twice with PBS, and then overlayed with a 1:1 mix of fresh and clarified conditioned CN medium supplemented with 25 µg/mL LP2 antibody. Virus spread was monitored by live-cell fluorescence imaging using an Incucyte SX5 (Sartorius), recording phase contrast images, green fluorescence and orange fluorescence every 3 h. Data were analysed using the Basic Analyser algorithm in the Incucyte analysis software (Sartorius). Half the volume of CN medium was replaced every 3 days.

## Results

### Optimisation of the i3Neurone differentiation protocol for infection studies

The protocols for iPSC culture and i3Neurone differentiation were adapted from [49] to increase the efficiency of differentiation and promote survival during maturation (Fig. 1a). Specifically, duration of the Accutase dissociation was decreased to reduce cell death. The plating procedure was modified by seeding cells in half the final culture volume of medium and incubating the cells on the plate for 15 min at RT before adding the remaining medium. This ensured neurones evenly coated the well and reduced clumping of neurone cell bodies. To ensure iPSCs had correctly differentiated into neurones following these adaptations, immunoblotting (Fig. 1b) and immunofluorescence microscopy (Fig. 1c) of cultured iPSCs and i3Neurones was performed. Immunoblotting confirmed the loss of pluripotency marker OCT4 and the gain of neuronal markers TAU and βIII tubulin following differentiation. The presence of extensive neurites and neuronal cytoskeletal proteins MAP2 and TAU (Fig. 1c) confirmed successful differentiation.

**Fig. 1.**
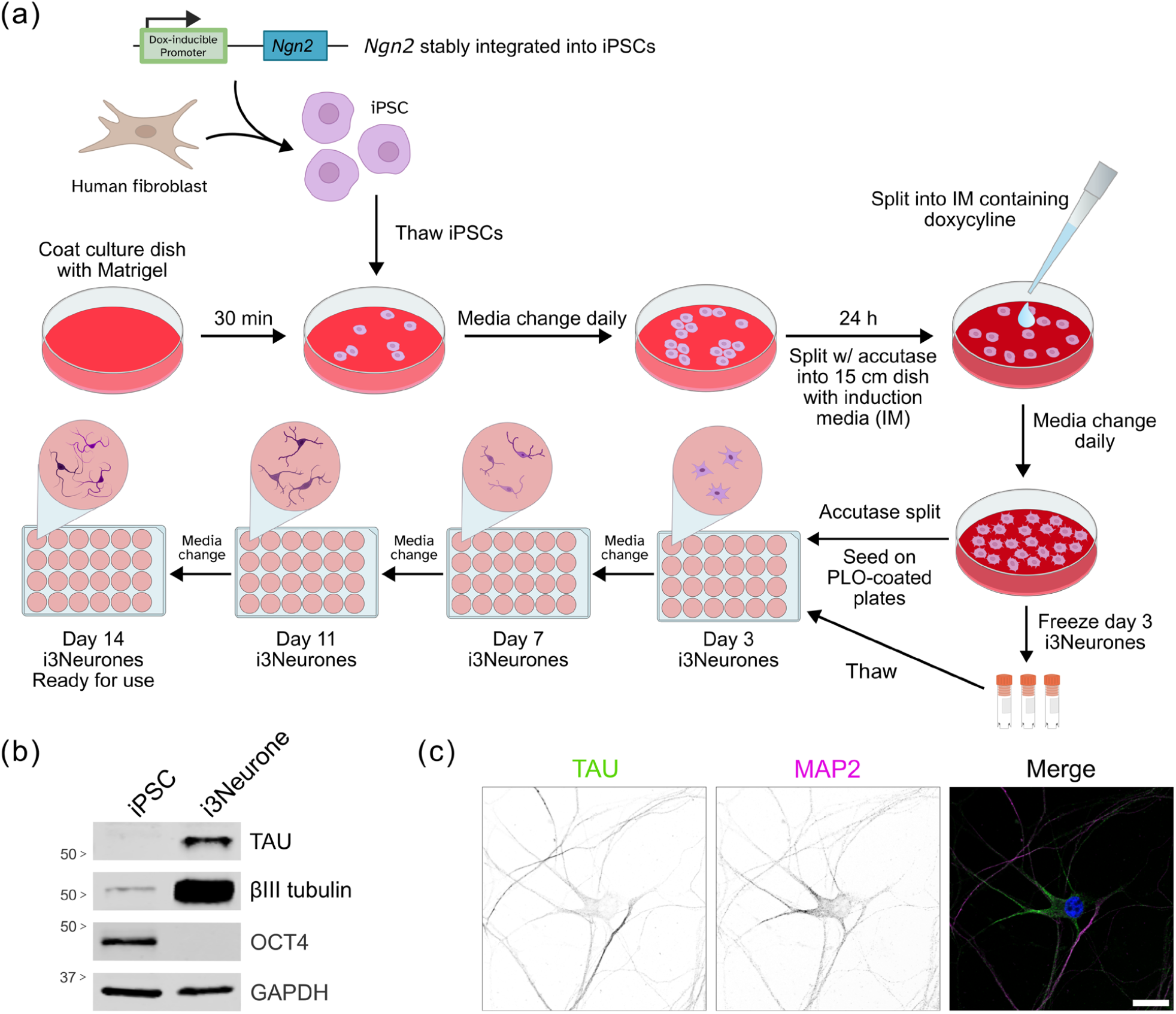
Differentiation of human iPSCs into cortical glutamatergic neurones (i3Neurones). (a) Schematic of the differentiation procedure. PLO, poly-L-ornithine; *Ngn2*, Neurogenin 2. (b) Validation of iPSC differentiation into neurones. Lysates of iPSCs and i3Neurones were immunoblotted for pluripotency marker OCT4 and neuronal markers TAU and βIII-tubulin. GAPDH is a loading control. (c) Confocal microscopy of differentiated i3Neurones, showing neuron-like morphology. Neuronal markers TAU (green) and MAP2 (magenta) are shown, and the merge image includes DAPI (blue). 20 µm scale bar.

### Optimisation of i3Neurone infection protocol

Initial infection tests suffered from technical issues including cell death, cell layers peeling off the plate and lower levels of infection than expected given the titre of virus inoculum. Several strategies were employed to combat these issues, testing different inoculation conditions. The first optimisation was to increase the volume of medium used for virus inoculation, as otherwise i3Neurones dried out and died during the inoculation step (Fig. S1a). Using approximately 30–50% of the final overlay volume for inoculation prevented such cell death. Secondly, to prevent peeling of the neurone layer (Fig. S1b) it was important to pipette liquids dropwise directly onto the cell layer rather than pipetting liquid onto the walls of the culture vessel [49].

To test if inoculation conditions could be optimised to increase the proportion of cells infected, neurones were infected at an MOI of either 1 or 5 via incubation with inoculum for 60, 90 or 120 min, either with continual rocking on a rocking platform or with manual rocking every 15 min (Fig. 2a). Cells were fixed at 16 hpi and stained for ICP4, an immediate-early viral protein that localises predominantly to the nucleus [65], plus propidium iodide to visualise nuclear DNA. Cells were imaged via automated microscopy to determine the percentage of infected cells (Fig. 2b). Automatic or manual rocking did not make a statistically significant difference to efficiency of infection at either MOI (two-way ANOVA of three independent experiments, p = 0.718 [MOI 1] or 0.280 [MOI 5]). At MOI 1 there was a significant increase in infection efficiency with increased duration of inoculation (Fig. 2a), but even after 2 h incubation with inoculum the proportion of infected cells remained below the theoretical maximum of 63.2% expected if infections occurred randomly. Extending the inoculum incubation time at MOI 5 did not alter the efficiency of infection, but use of an automated rocker resulted in significantly less variability of infection level across the replicates and time points (Levine’s test, p = 0.0346).

**Fig. 2.**
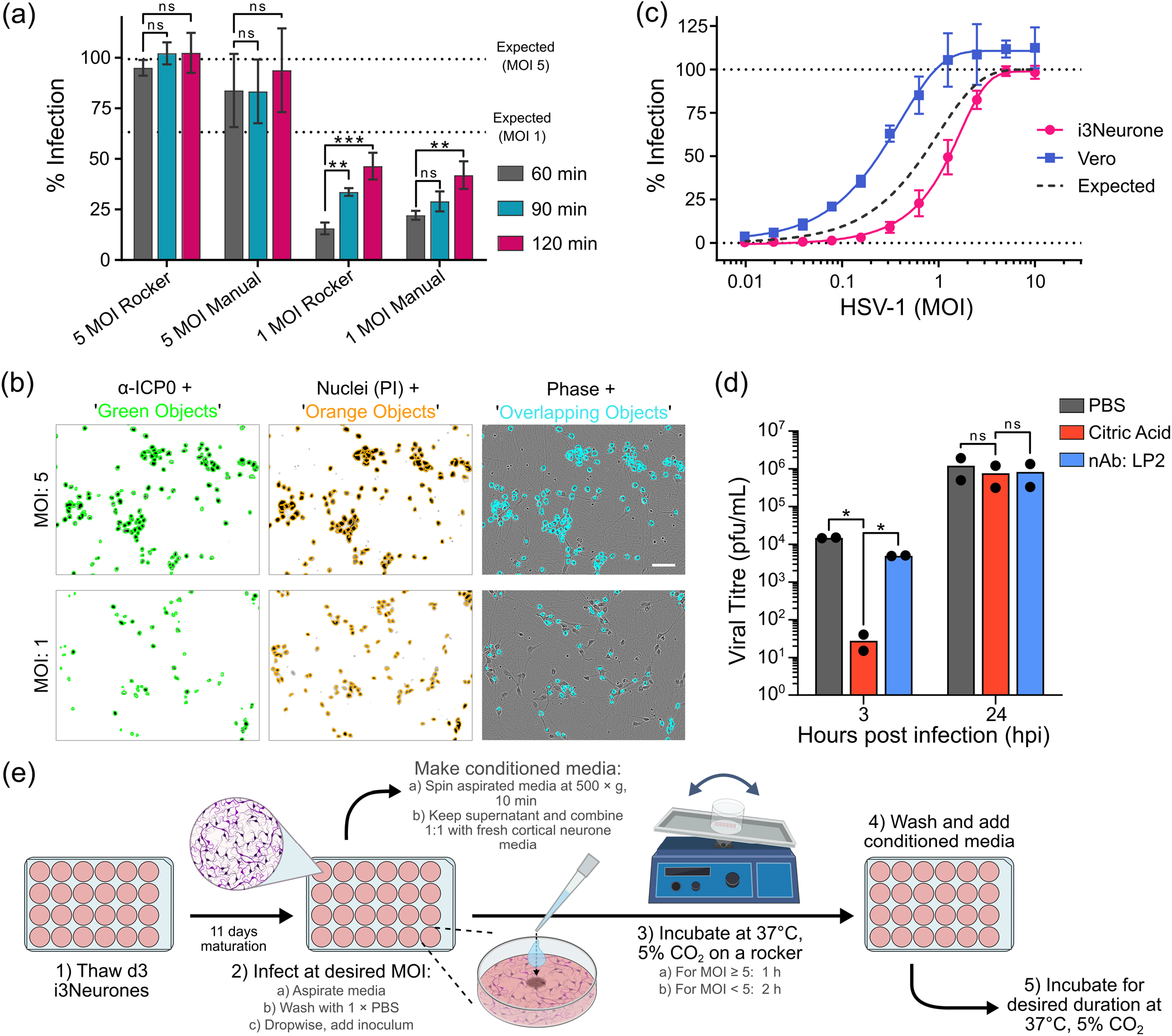
Optimisation of i3Neurone infection by HSV-1. (a) i3Neurones were infected at MOI 5 or 1 and incubated for the listed duration with manual or automated rocking. Percent of infected cells as determined by automated microscopy (mean ± SD from three independent experiments) is shown. Two-way ANOVA (automated or manual rocker versus time) confirmed no significant effect of rocking method but a significant effect of time for MOI 1 but not MOI 5. ns, no significance; **, p < 0.005; *** p < 0.001. (b) Representative automated microscopy images of infected i3Neurones. Objects where ICP4 (green, left) and propidium iodide (orange, middle) signals overlap (cyan, right) are counted as infected cells, expressed as a percentage of total propidium iodide objects (total cells). 50 μm scale bar. (c) Percentage of i3Neurones and Vero cells infected at different MOI (mean ± SD from three independent experiments). The percentage expected if infections occur randomly is shown. Scale bar represents 50 μm. (d) Optimisation of inoculum inactivation. Inoculum was removed at 1 hpi (MOI 5 HSV-1) and i3Neurones were either washed with PBS, incubated with neutralising antibody (nAb LP2) for 15 min, or washed with citric acid pH 3.0. Cells were harvested at 3 or 24 hpi and virus titres were determined by plaque assay. Mean and data points from two independent experiments, each performed in technical duplicate. One-way ANOVA confirms that the citrate wash yields a significant inactivation of input virus (3 hpi) but no difference in virus production at 24 hpi. ns, no significance; *, p < 0.05. (e) Schematic diagram of the optimised workflow for infecting i3Neurones with HSV-1.

To further investigate the efficiency of infection, a 2-fold dilution series of WT HSV-1 (from MOI 10 to 0.01) was used to inoculate i3Neurones and Vero cells in parallel (Fig. 2c). Below MOI 5 the proportion of infected cells was consistently lower than theoretically expected. Interestingly, a higher proportion of Vero cells were infected than expected, suggesting that virus titration via plaque assay may systematically underestimate the infectious titres. At high MOI, automated measurement of infection in Vero cells suggested that >100% of cells were infected. This arose due to the non-homogenous distribution of ICP4 staining in nuclei, with some infected Vero cell nuclei being counted twice by the Incucyte software (Fig. S2). Such double-counting may have also contributed to the greater-than-expected level of infection observed for Vero cells at different MOIs. Since double-counting of infected cells was not observed for i3Neurones, this phenomenon was not investigated further.

For temporally-resolved experiments like high MOI (single-step) growth curves that require a synchronous infection, it is necessary to inactivate any input virus particles that have not entered cells after a fixed time. This inactivation is often achieved via low pH treatment [66, 67]. Because the neurones may be more sensitive to chemical treatment than other cell lines, different methods were used to assess their inactivation efficiency whilst preserving cell viability. Differentiated i3Neurones were infected at MOI 5 using an automated plate rocker. After 1 h the inoculum was removed and cells were washed with citric acid, incubated with a potent neutralising antibody (LP2), or washed with PBS. The cultures were harvested at 3 and 24 hpi to assess both inoculum inactivation and subsequent virus production, which would decrease dramatically if cell viability was affected (Fig. 2d). The citric acid wash was the most effective treatment for inactivation, reducing the viral titre to below detection in some replicates. The antibody incubation was marginally more effective than PBS washing alone at removing input virus, but both were inferior to a citrate wash. There was no significant difference in virus yield at 24 hpi between any of the three conditions, demonstrating that none of these protocols adversely affected neurone viability. Based on these optimisations, refined protocols for low- and high-MOI infection of i3Neurones are summarised in Fig. 2e.

During the optimisation of the infection protocol, it was noted that neurones seemed highly tolerant of infection, with reduced morphology changes and prolonged survival compared to other cell types like Vero. The survival of i3Neurones following synchronous high MOI infection was thus investigated. Neurones were infected at MOI 5 with HSV-1 strains KOS, strain 17 or SC16, or mock-infected. Neurones were then incubated in the presence of propidium iodide, a DNA stain excluded from live cells, and imaged every 6 h via automated microscopy. By 5 days post-infection the neurones displayed morphology changes, with changes in the appearance of the soma and, in the case of KOS, clustering of the soma that is potentially indicative of syncytium formation. However, the neurones remained alive for upwards of 8 (KOS) or 10 (strain 17 and SC16) days, with visibly axonal degradation occurring close to the time of cell death (Fig. 3).

**Fig. 3.**
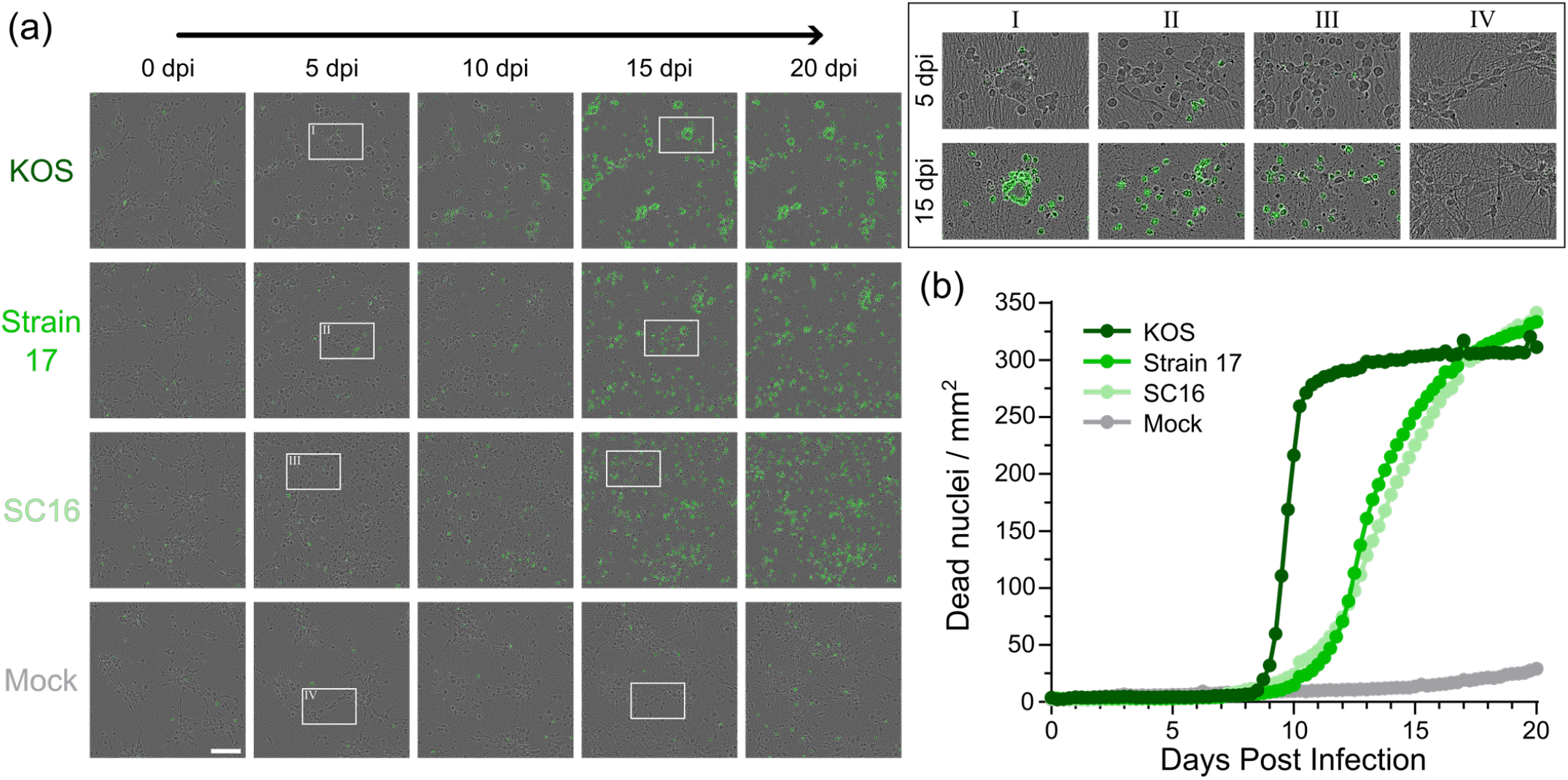
I3Neurones survive for over one week following HSV-1 infection. (a) Live-cell microscopy of i3Neurones infected at MOI 5 with HSV-1 strains KOS, strain 17 and SC16, at listed days post infection (dpi). Propidium iodide signal, which is excluded from live cells, is shown in green. Scale bar 100 µm. (b) Quantification of neurone survival, with cell death measured as increased number of propidium iodide positive nuclei (dead nuclei / mm^2^). Data are representative of three independent experiments.

### Optimised fixation and permeabilization of i3Neurones for immunocytochemistry

Neurites often became visibly damaged during the fixation and staining procedures used for preliminary infection quantification experiments, presumably due to their delicate nature. Different fixation conditions were thus tested for i3Neurones grown on coverslips that had been acid-etched to improve neurone adhesion [68]. Two conditions commonly used for immunocytochemistry of epithelial cells were tested (4% followed by 8% formaldehyde in 250 mM HEPES pH 7.4, or 100% methanol), as were two identified in the literature as being more effective for neurones or for preserving cytoskeletal structures (3% glyoxal pH 4.0 in 20% ethanol, or 4% formaldehyde in a cytoskeletal preservation solution)[69, 70]. Two different permeabilisation solutions were trialled, either 0.2% Triton X-100 for 5 min or 0.1% saponin for 15 min. i3Neurones were stained for the neuronal cytoskeletal protein β-III tubulin, the *trans*- Golgi network protein TGN46, the nuclear marker of infection ICP4, and for DNA using DAPI. The blocking and staining protocols used for each fixation/permeabilisation condition was identical, except for the inclusion of 0.01% saponin in the blocking buffer for cells permeabilised with saponin. Wide-field microscopy of coverslips fixed using the different protocols and permeabilised with Triton X-100 or saponin are shown in Figs S3 and S4, respectively.

Fixation using 4% then 8% formaldehyde in 250 mM HEPES yielded the best preservation of cells morphology, with excellent preservation of neurites. Triton permeabilization yielded visibly brighter signal for TGN46 and similar staining for the other markers, both in wide field (Figs S3 and S4) and confocal (Fig. 4) microscopy. Acquiring confocal Z-stacks (7–14 nm) proved the most reliable method for imaging both the thin neurites and the thicker cell bodies. It is notable that at 16 hpi there is no apparent change in overall cell morphology (contrast Fig. 1c and Fig. 4).

**Fig. 4.**
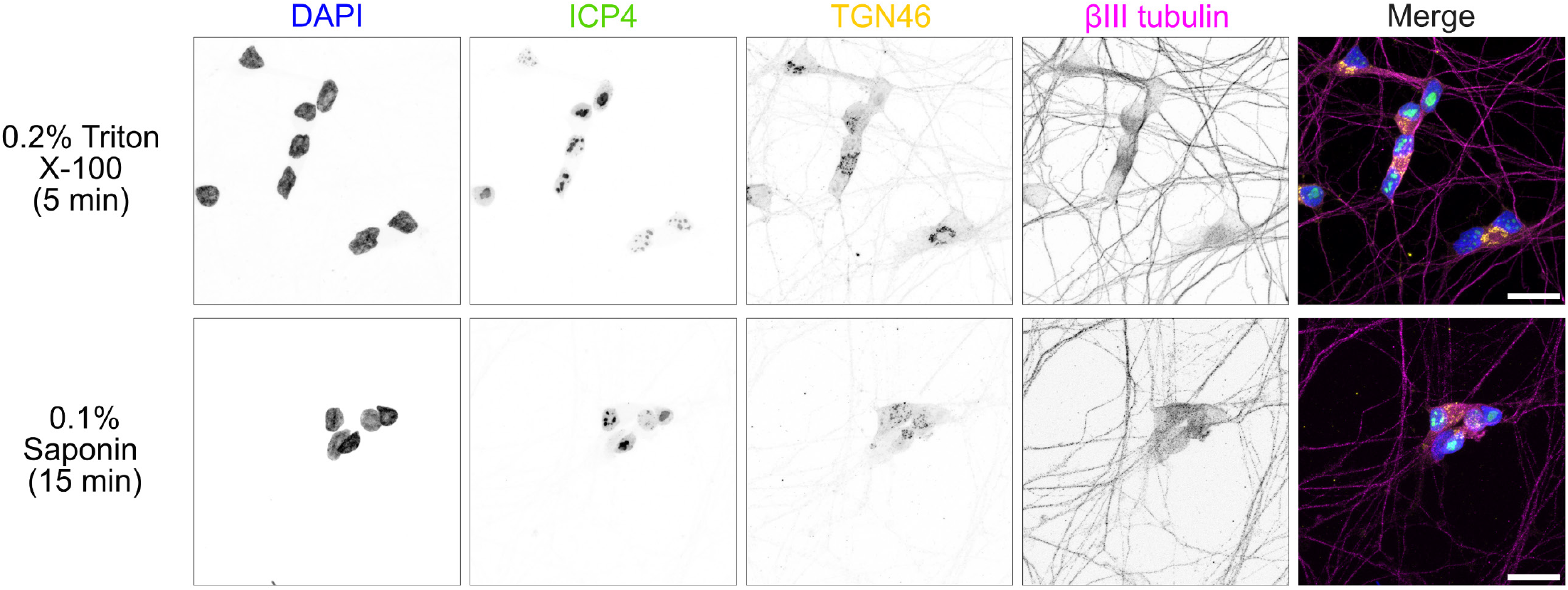
Confocal microscopy of HSV-1 infected (MOI 5) i3Neurons, fixed 16 hpi using 4% then 8% formaldehyde in 250 mM HEPES buffer and permeabilised using either 0.2% Triton X-100 for 5 min or 0.1% Saponin for 15 min. Cells were stained for ICP4 (green), TGN46 (yellow), βIII tubulin (magenta) and DNA (DAPI, blue) and maximum-intensity projections of 7 nm (Triton X-100) or 14 nm (saponin) Z stacks are shown. Triton X-100 permeabilisation yields stronger cytoskeletal (βIII tubulin) and organelle (TGN46) staining. 20 µm scale bar.

### i3Neurones as a model for HSV-1 lytic infection of cortical neurones

Having optimised the infection procedure, the utility of i3Neurones for monitoring HSV-1 spread was assessed. Plaque assays, which monitor the spread of virus to adjacent cells following infection of a single cell, are a well-established technique for measuring HSV-1 cell-to-cell spread in cells of the periphery like fibroblasts [30, 71]. However, the sparsity of neuronal cell bodies and potential for long-distance spread via intracellular transport of virions along neurites confounds the use of plaque assays to measure HSV-1 spread in i3Neurones. Therefore, virus neurone-to-neurone spread was monitored by infecting i3Neurones at low MOI (0.1) with HSV-1 strain KOS expressing the early protein ICP0 tagged with EYFP and the late protein gC tagged with mCherry [60]. Neutralising antibody (LP2) was included in the culture medium to inhibit cell-free spread and the spread of fluorescence, a proxy for infection spread, was monitored every 3 h by automated microscopy. While no signal was visible for ICP0-EYFP, a robust gC-mCherry signal was evident and this signal spread throughout the culture over the course of 96 h (Fig. 5a), confirming productive neurone-to-neurone HSV-1 spread. The infection appeared to spread most rapidly between the soma of adjacent cells but by 48 hpi infection was also evident in the soma of distant cells, consistent with spread of the infection via axonal transport of newly produced HSV-1 virions.

**Fig. 5.**
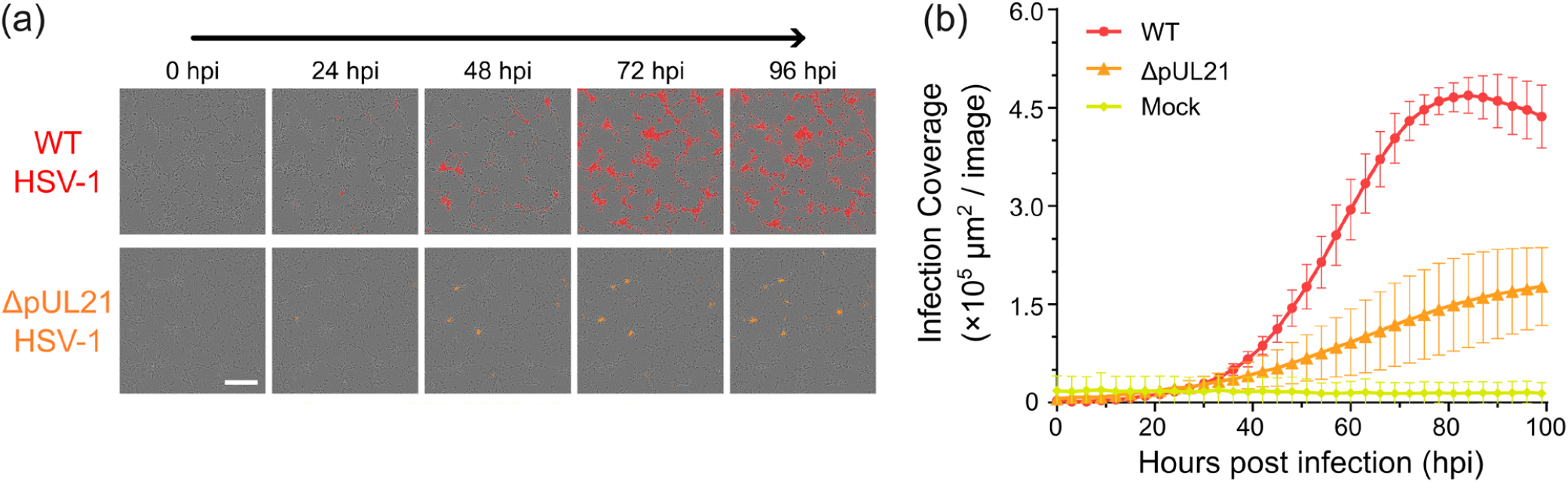
HSV-1 neurone-to-neurone spread. (a) Live-cell microscopy of i3Neurones infected at low MOI (0.1) with WT or ΔpUL21 timestamp HSV-1. gC-mCherry signal is shown in red or orange for WT and ΔpUL21 HSV-1, respectively. Scale bar 200 µm. (b) Quantification of timestamp virus spread, measured as increase area of gC-mCherry fluorescence (μm^2^ / image) for 4 days post infection. Mean ± SD for two independent experiments performed in technical triplicate are shown.

Many proteins present within the tegument layer of HSV-1 are known to contribute to efficient cell-to-cell spread in fibroblasts or keratinocytes [61, 72]. However, the contributions these proteins make to neurone-to-neurone spread is less clear. The role of tegument protein pUL21 is of particular interest, as its effect upon virus spread is known to vary by HSV strain and by cell type [73]. To assess its contribution to neurone-to-neurone spread, a mutant virus lacking expression of pUL21 was generated in the timestamp background (timestamp ΔpUL21, Fig. S5). The spread of timestamp ΔpUL21 infection was monitored via automated microscopy following low MOI (0.1) infection of i3Neurones (Fig. 5a). The spread of timestamp ΔpUL21 was substantially delayed when compared to the wild-type timestamp virus (Fig. 5b). However, there was still evidence of spread to soma of neurons distal to the initial site of infection, suggesting that transport of virions along neurites had not been completely impaired. Taken together, this experiment demonstrates how i3Neurones can be combined with fluorescently tagged HSV-1 to measure the effects of HSV-1 proteins on viral neurone-to-neurone spread.

In addition to monitoring virus spread, it is often desirable to monitor the abundance of cellular or viral proteins in synchronous populations of infected cells. While immunoblotting is a convenient technique for monitoring protein abundance, it can be difficult to obtain enough infected-cell lysate for immunoblotting when working with organoids or primary neurones. The scalability of i3Neurones [49], combined with the ability to perform efficient synchronous infection (Fig. 2), overcomes this limitation. To assess the feasibility of using immunoblots to monitor changes in host protein abundance, single wells of a 6-well dish containing 2×10^6^ i3Neurones were synchronously infected (MOI 5) with HSV-1 strains KOS, strain 17 and SC16. At 24 hpi cells were harvested, lysed and subjected to immunoblot analysis. For all three strains the viral capsid protein VP5 could be detected, confirming successful infection and late gene expression (Fig. 6). Additionally, compared to the mock-infected sample, all three infected lysates showed lower abundance of the cellular protein GOPC, a known target of HSV pUL56-mediated degradation [30]. This confirms that the i3Neurone system is suitable for biochemical analysis of HSV-1 neuronal infection.

**Fig. 6.**
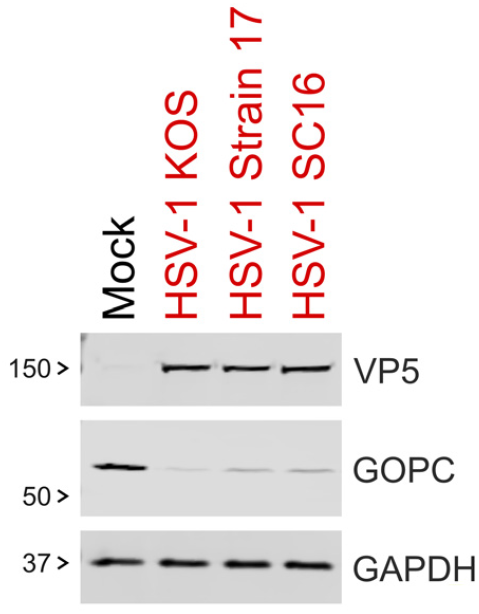
Validation of viral gene expression and function in i3Neurones by immunoblot. i3Neurones were infected at MOI 5 with indicated HSV-1 strains and lysed 24 hpi. Samples were immunoblotted for infection marker VP5, the cellular protein GOPC that is a target of pUL56-mediated degradation, and the cellular loading control GAPDH.

## Discussion

Here we present optimised protocols for the differentiation of human iPSC-derived cortical glutamatergic neurones (i3Neurones) and their infection with HSV-1. The i3Neurone system is highly scalable, allowing production of >10^7^ differentiated neurones with ease, and these neurones can be synchronously infected with high (>90%) efficiency (Fig. 2). These neurons survive for upwards of 8 days following infection (Fig. 3), consistent with previous reports of sympathetic mouse neurones surviving for up to 30 days following lytic infection with HSV-1 [74]. We show that i3Neurones are suitable for biochemical analysis of lytic HSV-1 infection (Fig. 6) and i3Neurones thus show strong potential for use in high resolution infection proteomics analysis [29, 30]. We have previously shown that i3Neurones can be infected with Zika virus [75], human astroviruses [76] and human enteroviruses [77]. i3Neurones thus represent a promising platform for advanced biochemical analysis of many neurotropic virus infections.

In addition to biochemical analyses, the reproducibility of i3Neurone differentiation [48] and their amenability to gene overexpression [51] or knockdown via the integrated dead Cas9 [50– 52] make them a powerful platform for functional analysis. We show here that i3Neurones can be combined with fluorescent virus strains of HSV-1 to monitor neurone-to-neurone spread of HSV-1. While we observed a signal for ICP0-EYFP in i3Neurones when imaged using a wide-field microscope, consistent with prior studies using timestamp HSV-1 [60, 78], we could not visualise ICP0-EYFP using the Incucyte SX5 automated microscope. This difference is likely to arise from a combination of lower ICP0 expression in i3Neurones, the use of a long working distance objective with low numerical aperture, plus suboptimal matching of the excitation (453–485 nm) and emission (494–533 nm) filters on our automated microscope to the EYFP fluorophore (peak excitation 515 nm and emission 530 nm). In the presence of neutralising antibody, we show that HSV-1 strain KOS spreads to the soma of neurones far from the initial site of infection within 48 hpi, consistent with intracellular transport of virions along neurites. It is unclear whether this spread represents virus particles budding from the soma of an infected cell, entering a neurite and undergoing retrograde transport to the nucleus, or whether it represents anterograde transport of newly assembled virions to neurite termini where they then bud to infect other neurones. Since HSV-1 strain KOS lacks a functional pUS9 protein [79], known to be important for both anterograde axonal transport and virus assembly at axon termini [80], it seems likely that the observed long-distance spread represents retrograde transport following infection of neurites. This could be confirmed in future studies using directional infection of soma or neurites in compartmentalised culture systems [81].

In summary, using HSV-1 as a model we have demonstrated the i3Neurone system to be a robust tool for measuring the replication and spread of viruses in cortical neurones. We anticipate that optimised neurone culture, infection and analysis protocols presented here will accelerate research into a broad range of clinically important neurotropic infections.

## Author contributions

Conceptualisation: JED, SCG; Funding Acquisition: JED, SCG; Investigation: DAN, ASN, HGB, VC; Project Administration: JED, SCG; Resources: CMC, JED; Supervision: JED, SCG; Visualisation: DAN; Writing – Original Draft Preparation: DAN, SCG; Writing – Review & Editing: DAN, HGB, AN, JED, SCG

## Acknowledgements

We thank Dr Michael Ward for the i3Neurones, Dr Gopal Sapkota for the Flp-In T-REx U2OS cells, Profs Stacey Efstathiou and Tony Minson for HSV-1 isolates, and the Cambridge Microscopy Bioscience Platform for their support and assistance in this work.

## References

1. Matthews E, Beckham JD, Piquet AL, Tyler KL, Chauhan L, et al. Herpesvirus-Associated Encephalitis: an Update. Curr Trop Med Rep 2022;9:92–100.

2. Jorgensen LK, Dalgaard LS, Ostergaard LJ, Norgaard M, Mogensen TH. Incidence and mortality of herpes simplex encephalitis in Denmark: A nationwide registry-based cohort study. J Infect 2017;74:42–49.

3. Ong KC, Wong KT. Understanding Enterovirus 71 Neuropathogenesis and Its Impact on Other Neurotropic Enteroviruses. Brain Pathol 2015;25:614–624.

4. Muñoz LS, Parra B, Pardo CA, Neuroviruses Emerging in the Americas Study. Neurological Implications of Zika Virus Infection in Adults. J Infect Dis 2017;216:S897–S905.

5. de Armas Fernández JR, Peña García CE, Acosta Herrera B, Betancourt Plaza I, Gutiérrez de la Cruz Y, et al. Report of an unusual association of Oropouche Fever with Guillain-Barré syndrome in Cuba, 2024. Eur J Clin Microbiol Infect Dis 2024;43:2233–2237.

6. Freitas DA, Souza-Santos R, Carvalho LMA, Barros WB, Neves LM, et al. Congenital Zika syndrome: A systematic review. PLoS One 2020;15:e0242367.

7. Marcocci ME, Napoletani G, Protto V, Kolesova O, Piacentini R, et al. Herpes Simplex Virus-1 in the Brain: The Dark Side of a Sneaky Infection. Trends Microbiol 2020;28:808–820.

8. Liu Y, Johnston C, Jarousse N, Fletcher SP, Iqbal S. Association between herpes simplex virus type 1 and the risk of Alzheimer’s disease: a retrospective case-control study. BMJ Open 2025;15:e093946.

9. Araya K, Watson R, Khanipov K, Golovko G, Taglialatela G. Increased risk of dementia associated with herpes simplex virus infections: Evidence from a retrospective cohort study using U.S. electronic health records. J Alzheimers Dis 2025;104:393–402.

10. Canova PN, Charron AJ, Leib DA. Models of Herpes Simplex Virus Latency. Viruses 2024;16:747.

11. Sun G, Viejo-Borbolla A. In Vitro Culture of Primary Mouse Neurons to Study Neuronal Infection. Methods Mol Biol 2025;2950:73–85.

12. Wilson AC. Impact of Cultured Neuron Models on α-Herpesvirus Latency Research. Viruses 2022;14:1209.

13. Wilcox CL, Johnson EM. Nerve growth factor deprivation results in the reactivation of latent herpes simplex virus in vitro. J Virol 1987;61:2311–2315.

14. Smith PR, Meyer A, Loerch S, Campbell ZT. Protocol for the isolation and culture of mouse dorsal root ganglion neurons for imaging applications. STAR Protoc 2023;4:102717.

15. Johansson PJ, Myhre EB, Blomberg J. Specificity of Fc receptors induced by herpes simplex virus type 1: comparison of immunoglobulin G from different animal species. J Virol 1985;56:489–494.

16. Verweij MC, Ressing ME, Knetsch W, Quinten E, Halenius A, et al. Inhibition of mouse TAP by immune evasion molecules encoded by non-murine herpesviruses. Mol Immunol 2011;48:835–845.

17. LaPaglia DM, Sapio MR, Burbelo PD, Thierry-Mieg J, Thierry-Mieg D, et al. RNA-Seq investigations of human post-mortem trigeminal ganglia. Cephalalgia 2018;38:912–932.

18. Rashidi AS, Tran DN, Peelen CR, van Gent M, Ouwendijk WJD, et al. Herpes simplex virus infection induces necroptosis of neurons and astrocytes in human fetal organotypic brain slice cultures. J Neuroinflammation 2024;21:38.

19. Lafaille FG, Pessach IM, Zhang S-Y, Ciancanelli MJ, Herman M, et al. Impaired intrinsic immunity to HSV-1 in human iPSC-derived TLR3-deficient CNS cells. Nature 2012;491:769–773.

20. Zimmer B, Ewaleifoh O, Harschnitz O, Lee Y-S, Peneau C, et al. Human iPSC-derived trigeminal neurons lack constitutive TLR3-dependent immunity that protects cortical neurons from HSV-1 infection. Proc Natl Acad Sci U S A 2018;115:E8775–E8782.

21. Dai Y, Idorn M, Serrero MC, Pan X, Thomsen EA, et al. TMEFF1 is a neuron-specific restriction factor for herpes simplex virus. Nature 2024;632:383–389.

22. Pourchet A, Modrek AS, Placantonakis DG, Mohr I, Wilson AC. Modeling HSV-1 Latency in Human Embryonic Stem Cell-Derived Neurons. Pathogens 2017;6:24.

23. Liu Z, Garcia Reino EJ, Harschnitz O, Guo H, Chan Y-H, et al. Encephalitis and poor neuronal death-mediated control of herpes simplex virus in human inherited RIPK3 deficiency. Sci Immunol 2023;8:eade2860.

24. D’Aiuto L, Bloom DC, Naciri JN, Smith A, Edwards TG, et al. Modeling Herpes Simplex Virus 1 Infections in Human Central Nervous System Neuronal Cells Using Two- and Three-Dimensional Cultures Derived from Induced Pluripotent Stem Cells. J Virol 2019;93:e00111–19.

25. Krenn V, Bosone C, Burkard TR, Spanier J, Kalinke U, et al. Organoid modeling of Zika and herpes simplex virus 1 infections reveals virus-specific responses leading to microcephaly. Cell Stem Cell 2021;28:1362-1379.e7.

26. Qiao H, Guo M, Shang J, Zhao W, Wang Z, et al. Herpes simplex virus type 1 infection leads to neurodevelopmental disorder-associated neuropathological changes. PLoS Pathog 2020;16:e1008899.

27. Cairns DM, Rouleau N, Parker RN, Walsh KG, Gehrke L, et al. A 3D human brain-like tissue model of herpes-induced Alzheimer’s disease. Sci Adv 2020;6:eaay8828.

28. Rybak-Wolf A, Wyler E, Pentimalli TM, Legnini I, Oliveras Martinez A, et al. Modelling viral encephalitis caused by herpes simplex virus 1 infection in cerebral organoids. Nat Microbiol 2023;8:1252–1266.

29. Fletcher-Etherington A, Weekes MP. Quantitative Temporal Viromics. Annu Rev Virol 2021;8:159–181.

30. Soh TK, Davies CTR, Muenzner J, Hunter LM, Barrow HG, et al. Temporal Proteomic Analysis of Herpes Simplex Virus 1 Infection Reveals Cell-Surface Remodeling via pUL56-Mediated GOPC Degradation. Cell Rep 2020;33:108235.

31. Weekes MP, Tomasec P, Huttlin EL, Fielding CA, Nusinow D, et al. Quantitative temporal viromics: an approach to investigate host-pathogen interaction. Cell 2014;157:1460–1472.

32. Shipley MM, Mangold CA, Kuny CV, Szpara ML. Differentiated Human SH-SY5Y Cells Provide a Reductionist Model of Herpes Simplex Virus 1 Neurotropism. J Virol 2017;91:e00958–17.

33. Kang W, Mukerjee R, Fraser NW. Establishment and maintenance of HSV latent infection is mediated through correct splicing of the LAT primary transcript. Virology 2003;312:233–244.

34. Krishna A, Biryukov M, Trefois C, Antony PMA, Hussong R, et al. Systems genomics evaluation of the SH-SY5Y neuroblastoma cell line as a model for Parkinson’s disease. BMC Genomics 2014;15:1154.

35. Do JH, Kim IS, Park T-K, Choi D-K. Genome-wide examination of chromosomal aberrations in neuroblastoma SH-SY5Y cells by array-based comparative genomic hybridization. Mol Cells 2007;24:105–112.

36. Kovalevich J, Langford D. Considerations for the use of SH-SY5Y neuroblastoma cells in neurobiology. Methods Mol Biol 2013;1078:9–21.

37. Edwards TG, Bloom DC. Lund Human Mesencephalic (LUHMES) Neuronal Cell Line Supports Herpes Simplex Virus 1 Latency In Vitro. J Virol 2019;93:e02210–18.

38. Whisnant AW, Dyck Dionisi O, Salazar Sanchez V, Rappold JM, Djakovic L, et al. Herpes simplex virus 1 inhibits phosphorylation of RNA polymerase II CTD serine-7. J Virol 2024;98:e0117824.

39. Tüshaus J, Kataka ES, Zaucha J, Frishman D, Müller SA, et al. Neuronal Differentiation of LUHMES Cells Induces Substantial Changes of the Proteome. Proteomics 2021;21:e2000174.

40. Lauter G, Coschiera A, Yoshihara M, Sugiaman-Trapman D, Ezer S, et al. Differentiation of ciliated human midbrain-derived LUHMES neurons. J Cell Sci 2020;133:jcs249789.

41. Sili U, Kaya A, Mert A, HSV Encephalitis Study Group. Herpes simplex virus encephalitis: clinical manifestations, diagnosis and outcome in 106 adult patients. J Clin Virol 2014;60:112–118.

42. Cho H, Proll SC, Szretter KJ, Katze MG, Gale M, et al. Differential innate immune response programs in neuronal subtypes determine susceptibility to infection in the brain by positive-stranded RNA viruses. Nat Med 2013;19:458–464.

43. Ng AHM, Khoshakhlagh P, Rojo Arias JE, Pasquini G, Wang K, et al. A comprehensive library of human transcription factors for cell fate engineering. Nat Biotechnol 2021;39:510–519.

44. Deng Y, Lin Y, Chen S, Xiang Y, Chen H, et al. Neuronal miR-9 promotes HSV-1 epigenetic silencing and latency by repressing Oct-1 and Onecut family genes. Nat Commun 2024;15:1991.

45. Sun B, Yang X, Hou F, Yu X, Wang Q, et al. Regulation of host and virus genes by neuronal miR-138 favours herpes simplex virus 1 latency. Nat Microbiol 2021;6:682–696.

46. Oh HS, Chou S-F, Raja P, Shim J, Das B, et al. Validation of human sensory neurons derived from inducible pluripotent stem cells as a model for latent infection and reactivation by herpes simplex virus1. mBio 2025;e0187125.

47. Zhang Y, Pak C, Han Y, Ahlenius H, Zhang Z, et al. Rapid single-step induction of functional neurons from human pluripotent stem cells. Neuron 2013;78:785–798.

48. Wang C, Ward ME, Chen R, Liu K, Tracy TE, et al. Scalable Production of iPSC-Derived Human Neurons to Identify Tau-Lowering Compounds by High-Content Screening. Stem Cell Reports 2017;9:1221–1233.

49. Fernandopulle MS, Prestil R, Grunseich C, Wang C, Gan L, et al. Transcription Factor-Mediated Differentiation of Human iPSCs into Neurons. Curr Protoc Cell Biol 2018;79:e51.

50. Nicholson AS, Priestman DA, Antrobus R, Williamson JC, Bush R, et al. Plasma membrane remodeling in GM2 gangliosidoses drives synaptic dysfunction. PLoS Biol 2025;23:e3003265.

51. Rodger C, Flex E, Allison RJ, Sanchis-Juan A, Hasenahuer MA, et al. De Novo VPS4A Mutations Cause Multisystem Disease with Abnormal Neurodevelopment. Am J Hum Genet 2020;107:1129–1148.

52. Tian R, Gachechiladze MA, Ludwig CH, Laurie MT, Hong JY, et al. CRISPR Interference-Based Platform for Multimodal Genetic Screens in Human iPSC-Derived Neurons. Neuron 2019;104:239-255.e12.

53. Kim DI, Jensen SC, Noble KA, Kc B, Roux KH, et al. An improved smaller biotin ligase for BioID proximity labeling. Mol Biol Cell 2016;27:1188–96.

54. Bozatzi P, Dingwell KS, Wu KZ, Cooper F, Cummins TD, et al. PAWS1 controls Wnt signalling through association with casein kinase 1α. EMBO Rep 2018;19:e44807.

55. Monkhouse H, Carter-Lopez DS, Benedyk TH, Deane JE, Graham SC. Alphaherpesvirus pUL21 homologues use non-canonical motifs to compete with cellular adaptors for protein phosphatase 1 binding. BioRχiv online preprint. Epub ahead of print 25 July 2025. DOI: 10.1101/2025.07.22.666160.

56. Minson AC, Hodgman TC, Digard P, Hancock DC, Bell SE, et al. An analysis of the biological properties of monoclonal antibodies against glycoprotein D of herpes simplex virus and identification of amino acid substitutions that confer resistance to neutralization. J Gen Virol 1986;67:1001–1013.

57. McClelland DA, Aitken JD, Bhella D, McNab D, Mitchell J, et al. pH reduction as a trigger for dissociation of herpes simplex virus type 1 scaffolds. J Virol 2002;76:7407–7417.

58. Benedyk TH, Muenzner J, Connor V, Han Y, Brown K, et al. pUL21 is a viral phosphatase adaptor that promotes herpes simplex virus replication and spread. PLoS Pathog 2021;17:e1009824.

59. Tischer BK, von Einem J, Kaufer B, Osterrieder N. Two-step red-mediated recombination for versatile high-efficiency markerless DNA manipulation in Escherichia coli. Biotechniques 2006;40:191–197.

60. Scherer KM, Manton JD, Soh TK, Mascheroni L, Connor V, et al. A fluorescent reporter system enables spatiotemporal analysis of host cell modification during herpes simplex virus-1 replication. J Biol Chem 2021;296:100236.

61. Nahas KL, Connor V, Wijesinghe KJ, Barrow HG, Dobbie IM, et al. Applying 3D correlative structured illumination microscopy and X-ray tomography to characterise herpes simplex virus-1 morphogenesis. eLife reviewed preprint. Epub ahead of print 25 February 2025. DOI: 10.7554/eLife.105209.1.

62. Schindelin J, Arganda-Carreras I, Frise E, Kaynig V, Longair M, et al. Fiji: an open-source platform for biological-image analysis. Nat Methods 2012;9:676–682.

63. Rueden CT, Schindelin J, Hiner MC, DeZonia BE, Walter AE, et al. ImageJ2: ImageJ for the next generation of scientific image data. BMC Bioinformatics 2017;18:529.

64. Brown MB, Forsythe AB. Robust Tests for the Equality of Variances. Journal of the American Statistical Association 1974;69:364–367.

65. Zhu Z, Schaffer PA. Intracellular localization of the herpes simplex virus type 1 major transcriptional regulatory protein, ICP4, is affected by ICP27. J Virol 1995;69:49–59.

66. Huang AS, Wagner RR. PENETRATION OF HERPES SIMPLEX VIRUS INTO HUMAN EPIDERMOID CELLS. Proc Soc Exp Biol Med 1964;116:863–869.

67. MacLean CA. HSV Entry and Spread. Methods Mol Med 1998;10:9–18.

68. Sheta R, Teixeira M, Idi W, Oueslati A. Optimized protocol for the generation of functional human induced-pluripotent-stem-cell-derived dopaminergic neurons. STAR Protoc 2023;4:102486.

69. Stalder D, Yakunin I, Pereira C, Eden J, Gershlick DC. Recruitment of PI4KIIIβ to the Golgi by ACBD3 is dependent on an upstream pathway of a SNARE complex and golgins. Mol Biol Cell 2024;35:ar20.

70. Richter KN, Revelo NH, Seitz KJ, Helm MS, Sarkar D, et al. Glyoxal as an alternative fixative to formaldehyde in immunostaining and super-resolution microscopy. EMBO J 2018;37:139–159.

71. Albecka A, Owen DJ, Ivanova L, Brun J, Liman R, et al. Dual Function of the pUL7-pUL51 Tegument Protein Complex in Herpes Simplex Virus 1 Infection. J Virol 2017;91:e02196–16.

72. Ahmad I, Wilson DW. HSV-1 Cytoplasmic Envelopment and Egress. Int J Mol Sci 2020;21:E5969.

73. Finnen RL, Banfield BW. CRISPR/Cas9 Mutagenesis of UL21 in Multiple Strains of Herpes Simplex Virus Reveals Differential Requirements for pUL21 in Viral Replication. Viruses 2018;10:258.

74. Dochnal S, Merchant HY, Schinlever AR, Babnis A, Depledge DP, et al. DLK-Dependent Biphasic Reactivation of Herpes Simplex Virus Latency Established in the Absence of Antivirals. J Virol 2022;96:e0050822.

75. Lefèvre C, Cook GM, Dinan AM, Torii S, Stewart H, et al. Zika viruses encode 5’ upstream open reading frames affecting infection of human brain cells. Nat Commun 2024;15:8822.

76. Ali H, Lulla A, Nicholson AS, Hankinson J, Wignall-Fleming EB, et al. Attenuation hotspots in neurotropic human astroviruses. PLoS Biol 2023;21:e3001815.

77. O’Connor RL, Cook GM, Hankinson J, Fominykh K, Cheng SH, et al. Flexibility and modulation of translation initiation in enterovirus genomes. BioRχiv online preprint. Epub ahead of print 24 March 2025. DOI: 10.1101/2025.03.24.645098.

78. Nahas KL, Connor V, Scherer KM, Kaminski CF, Harkiolaki M, et al. Near-native state imaging by cryo-soft-X-ray tomography reveals remodelling of multiple cellular organelles during HSV-1 infection. PLoS Pathog 2022;18:e1010629.

79. Negatsch A, Mettenleiter TC, Fuchs W. Herpes simplex virus type 1 strain KOS carries a defective US9 and a mutated US8A gene. J Gen Virol 2011;92:167–172.

80. Miranda-Saksena M, Boadle RA, Diefenbach RJ, Cunningham AL. Dual Role of Herpes Simplex Virus 1 pUS9 in Virus Anterograde Axonal Transport and Final Assembly in Growth Cones in Distal Axons. J Virol 2015;90:2653–2663.

81. Tierney WM, Vicino IA, Sun SY, Chiu W, Engel EA, et al. Methods and Applications of Campenot Trichamber Neuronal Cultures for the Study of Neuroinvasive Viruses. Methods Mol Biol 2022;2431:181–206.

