## supplementary for "Optimisation of lytic herpes simplex virus infection in human induced pluripotent stem cell derived cortical neurones"

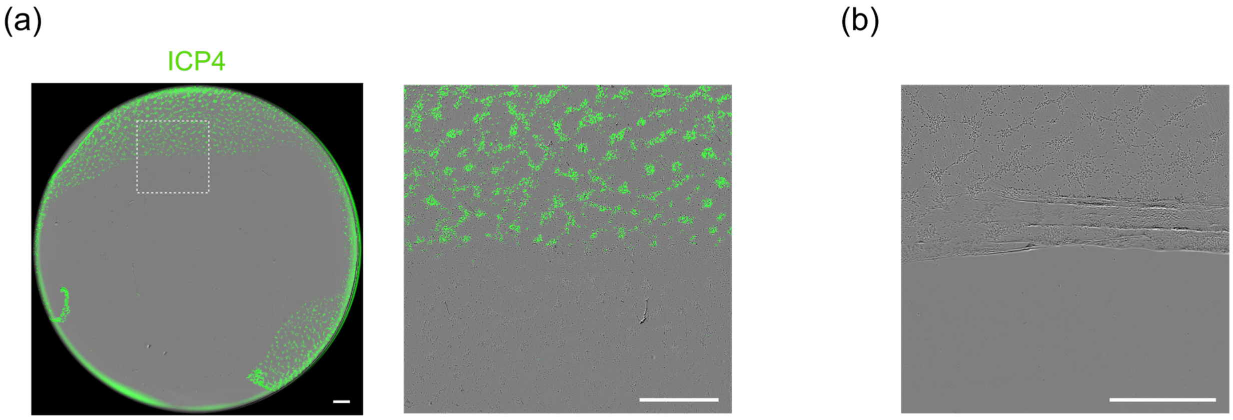


**Fig. S1.**

Neurone layers are prone to drying out and folding. (a) i3Neurones in a 24-well plate were infected at 5 MOI for 1 h in an inoculation volume of 200 µL before three PBS washes. 16 hpi cells were fixed, stained for ICP4 and imaged via whole-well scanning in an Incucyte SX5. The presence of fluorescence at the edges of the well was an indication of extensive cell death, due to the centre of the well drying out during virus inoculation. (b) Phase contrast image of i3Neurone layer peeling because of pipetting down the wall of the well. Scale bars 800 µm.

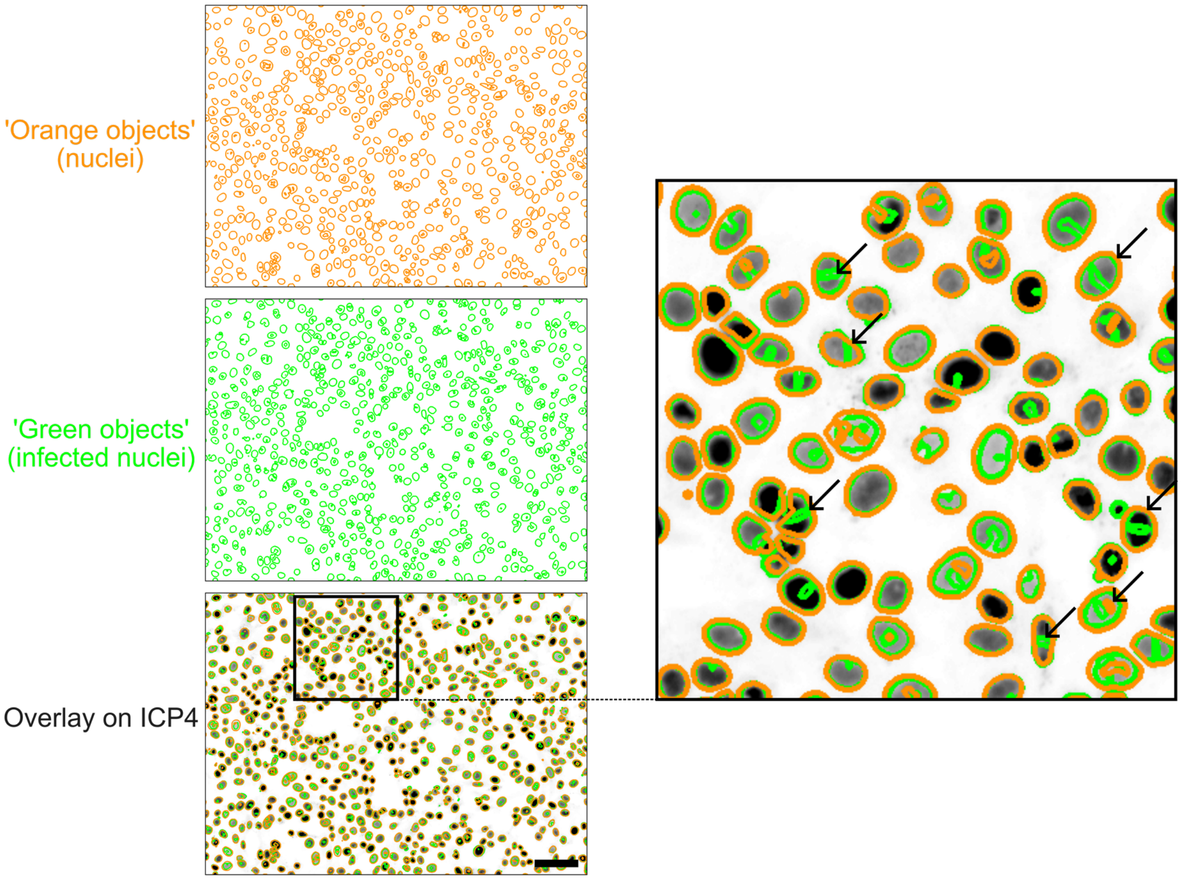


**Fig. S2.**

Incucyte SX5 ‘basic analyzer’ software double-counts infected Vero cells. Arrows indicate incidences where there are two distinct objects for ICP4 signal (green) within a single nucleus (orange), which inflates the number of infected nuclei. Scale bar 100 µm.

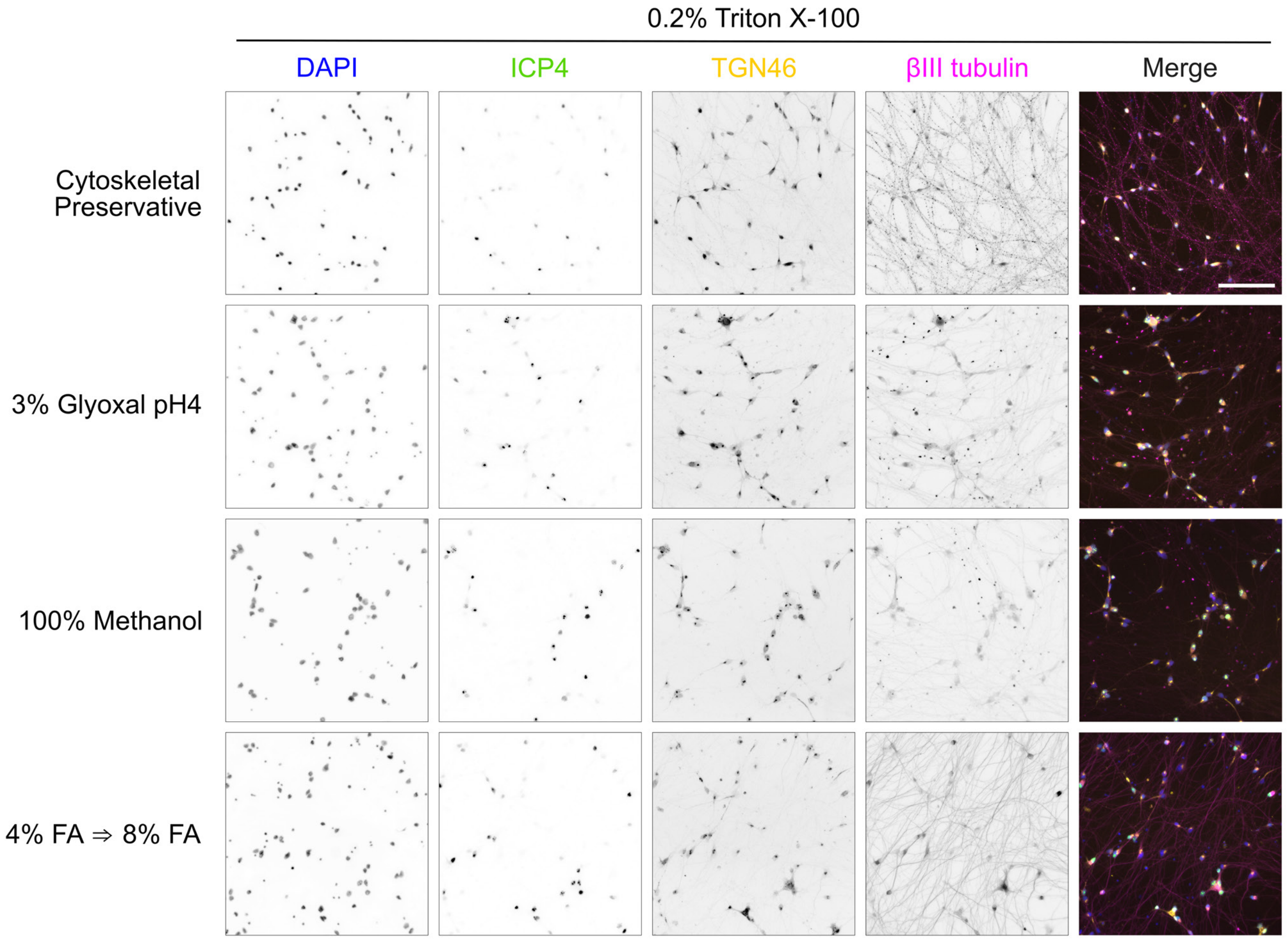


**Fig. S3.**

Exploration of different fixation protocols, showing that stepwise formaldehyde (FA) fixation gives the best preservation of neurone morphology and subcellular structures. I3Neurones were infected with HSV-1 (MOI 5) and fixed 16 hpi using the four listed fixatives, as detailed in the Methods. Fixed neurones were permeabilised with 0.2% Triton X-100 in PBS and immunostained for ICP4 (green), TGN46 (yellow) and βIII tubulin (magenta). During coverslip mounting the cells were also stained with DAPI (blue). Scale bar 100 µm.

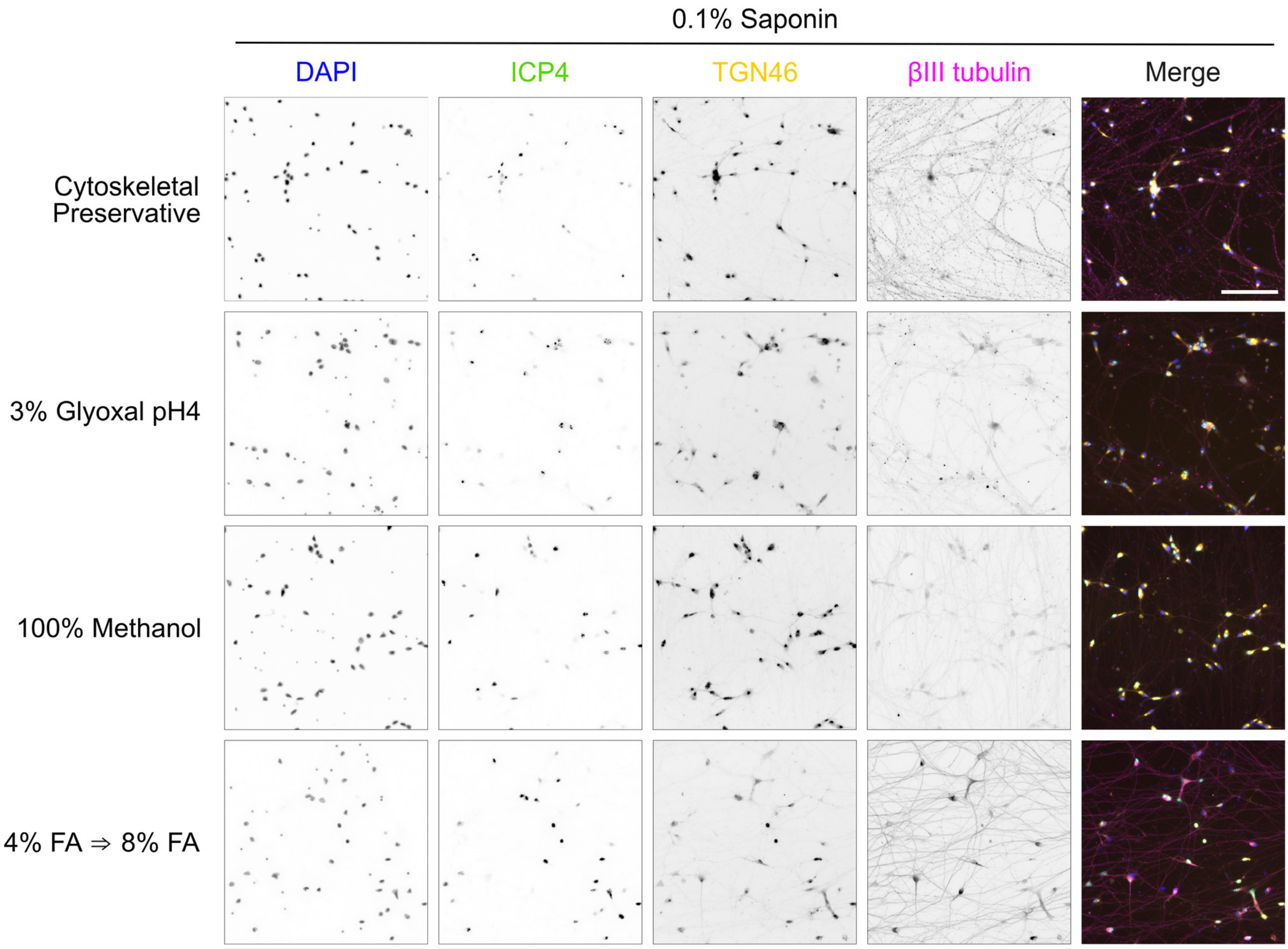


**Fig. S4.**

Exploration of different fixation protocols, showing that stepwise formaldehyde (FA) fixation gives the best preservation of neurone morphology and subcellular structures. I3Neurones were infected with HSV-1 (MOI 5) and fixed 16 hpi using the four listed fixatives, as detailed in the Methods. Fixed neurones were permeabilised with 0.1% saponin in PBS and immunostained for ICP4 (green), TGN46 (yellow) and βIII tubulin (magenta). During coverslip mounting the cells were also stained with DAPI (blue). Scale bar 100 µm.

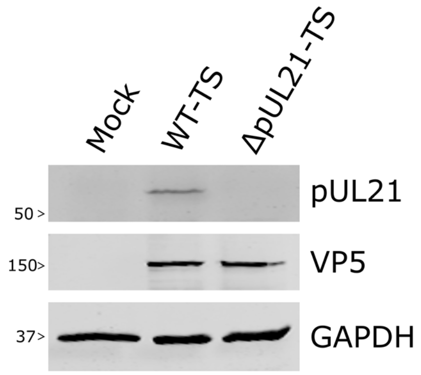


**Fig. S5.**

Timestamp ΔpUL21 HSV-1 lacks expression of viral protein pUL21. U2OS cells were infected at MOI 5 with WT and ΔpUL21 timestamp (TS) viruses, or mock-infected, and lysates harvested at 16 hpi. Lysates were immunoblotted for viral proteins pUL21 and VP5 plus the cellular protein GAPDH.
